# How frequently are insects wounded in the wild? A case study using *Drosophila melanogaster*

**DOI:** 10.1101/2023.08.25.554863

**Authors:** Bengisu S. Subasi, Veit Grabe, Martin Kaltenpoth, Jens Rolff, Sophie A.O. Armitage

## Abstract

1. Wounding occurs across multicellular organisms. There is a deep understanding of infections and immune responses from lab studies, yet wounds remain relatively comparatively understudied in nature. Ecological interactions like predator attacks, intra- and inter-specific competition can lead to wounding. Furthermore, rates of wounding may vary depending on factors such as sex and geographic location, with consequences for host mobility, reproduction, and susceptibility to pathogens.
2. Wounds initiate an immune response, resulting in the deposition of the brown-black pigment melanin in insects, and they are potential entry points for pathogens. Despite the potential ecological and evolutionary implications of wounding, and the relative abundance of lab immunity studies utilising *Drosophila melanogaster*, wounding in the wild in this model is unstudied. Our aim, therefore, was to investigate the prevalence and examine potential causes of wounds in wild-collected *D. melanogaster*.
3. From systematic collections of female and male flies, over three seasons and locations, we found that 31% of *D. melanogaster* were wounded. The abdomen was more frequently wounded than other body parts, and females were more likely to be injured particularly on the ventral abdomen, compared to males. Encapsulated parasitoid egg frequency was just under ten percent. Moreover, just under one percent of Drosophilidae species had mites attached to their body, the majority of which also caused wounds, i.e., were potentially parasitic.
4. Wounding is prevalent in *D. melanogaster*, and as such it is likely to exert selection pressure on host immunity for two reasons: on a rapid and efficient wound repair, and on responding to opportunistic infections. Wounds are thus expected to be important drivers of immune system evolution and to affect individual fitness and population dynamics.

## 1. Introduction

Wild organisms commonly incur wounds (Lindsay, 2010; Rennolds & Bely, 2023). Amongst others, wounds have been documented in wild vertebrates such as lizards (Fenner et al., 2008) and snakes (Willis et al., 1982), and invertebrates such as crustaceans (Plaistow et al., 2003), benthic invertebrates (Lindsay, 2010), and insects (Shapiro, 1974; Wallin, 1988; Cherrill & Brown, 1997). In insects, types of wounds include wing damage (Burkhard et al., 2002; Combes et al., 2010; Foster & Cartar, 2011; Rajabi et al., 2020), loss of antennae, legs, or bristles (Frank et al., 2018; Gilad et al., 2022), abdominal wounds (Cherrill & Brown, 1997) and copulatory wounds (Kamimura, 2007; Reinhardt et al., 2015). For example, around 30 % of adult bush crickets, *Decticus verrucivorus*, had antennal or leg damage (Cherrill & Brown, 1997). Even though there are many reports of wounds in wild insects, there is a paucity of studies that have systematically surveyed wounding in the wild (but see e.g., Gilad et al., 2022). Wounds can be important ecological and evolutionary factors, given that they can entail fitness costs and provide a portal for pathogen entry, but to estimate the impact of wounding systematic quantitative data is required on their occurrence.

We here focus on wounds in insects, which can occur for a variety of reasons, including predator attacks (Morin, 1985; Walters & Pawlik, 2005; Lindsay, 2010; Mukherjee & Heithaus, 2013; Frank et al., 2018; Reimchen & Bergstrom, 2023), intra- and inter-specific competition over food, territory, or mating (Stoks, 1998; Reinhardt et al., 2015, Liu et al., 2017) or wear and tear (Wallin, 1988) caused by environmental factors. For example, ants are wounded by termite soldiers and often lose limbs (Frank et al., 2018), and male monarch butterflies receive wing damage during mating (Leong et al., 1993). In some species, intersexual conflict results in copulatory wounds particularly to the females (Reinhardt et al., 2015; Dougherty et al., 2017). Lastly, parasites such as mites can wound the host cuticle with their mouth parts (chelicerae) when feeding on haemolymph (Kanbar & Engels, 2003).

Laboratory studies have shown that wounds can be costly. In severe cases they result in death (Gilad et al., 2022). They can negatively affect behaviour and cause significant changes in host physiology (Cartar, 1992; Carey et al., 2007; Leech et al., 2017; Rennolds & Bely, 2023). Wounded animals might be more susceptible to attack by predators (Brower, 1988; Harris, 1989) or parasites (Frank et al., 2018), and wounds can negatively affect reproduction (Harwood et al., 2013; Sepulveda et al., 2008; Shandilya et al., 2018; Von Wyschetzki et al., 2016). Wounds can have ecological consequences for predator-prey dynamics, population dynamics, and competitive interactions, and they can have evolutionary consequences through their effect on fitness and the selection pressure they impose on immune defences (Plaistow et al., 2003; Rennolds & Bely, 2023).

Importantly, wounds can be entry points for infections. Organisms are constantly in contact with the microbes in their living environment and the insect cuticle can harbour microbial communities (Ren et al., 2007; Birer et al., 2020). Once the cuticle is breached because of a wound, an opportunistic pathogen could potentially enter the body. For example, in a non-sterile environment the mortality of wounded ants was 80 % in 24 hours while it was only 10 % in a sterile environment (Frank et al., 2018). After a leg wound in *D. melanogaster* the spread of the bacteria into a fly body and the pathogenicity of the bacteria affected fly survival (Kari et al., 2013). Furthermore, parasitic mites can act as disease vectors in honeybees and increase the hosts’ susceptibility to viral, bacterial, and fungal infections (Glinski & Jarosz, 1992; Brødsgaard et al., 2000; Kanbar & Engels, 2003). Interestingly, mites have been experimentally shown to transmit *Spiroplasma poulsonii*, a male-killing endosymbiont of *Drosophila nebuosa* and *Drosophila willistoni*, from infected to uninfected flies, both within and between different species of *Drosophila* (Jaenike et al., 2007).

Wound healing is found across the animal kingdom (Arenas Gómez et al., 2020). In insects, wounds induce an immune response similar to that used to encapsulate parasites and pathogens. It has long been recognised that wounding in insects can result in brown to black melanised marks on the cuticle, for example topical scratching and abrasion induced by conspecifics in the silkworm *Bombyx mori* (Pasteur 1870, referenced in Brey & Hultmark, 1998) and experimentally scratched *D. melanogaster* larval cuticle (Önfelt Tingvall et al., 2001). Melanin is a pigment that plays a central role in insect immunity and cuticular darkening (Whitten & Coates, 2017). The production of melanin relies in part on phenoloxidase (PO) and its precursor prophenoloxidase (proPO), and proPO and its activating cascade have been detected in the cuticle of *B. mori* (Ashida & Brey, 1995). If the wound is more severe than a scratch, and the epidermis and basement membrane underlying soft cuticle are breached, haemolymph coagulation and clotting rapidly occur, thus preventing both haemolymph loss and microorganisms from passing into the haemocoel (Bidla et al., 2005; Dushay, 2009; Dziedziech et al., 2020). The proPO cascade is also immediately activated leading to melanisation and a hard clot (Dushay, 2009), which is observable as brown to black pigmentation. Importantly melanin can also be deposited on the surface of invading bacteria and fungi, thereby aggregating, and immobilising microbes (Zhao et al., 2007), as well as killing them through the production of cytotoxic side-products of the proPO cascade (Zhao et al., 2007; Nappi & Christensen, 2005). It is thought that coagulation and clotting do not occur in insects with a hard cuticle, because their haemolymph is under less than atmospheric pressure (Dushay, 2009); nonetheless wounds in adults are still melanised (e.g., Tang, 2009). The melanised wounds from within a larval instar or within the pupal or adult phase remain visible on the cuticle, and as such, they are a signature of wounding during that phase. Endoparasitoid wasps lay their eggs on or in the *Drosophila* body in the early host life history stages (larvae, or pupae; Godfray, 1994). The presence of wasp eggs triggers a haemocyte-mediated encapsulation reaction and if the immune response is successful the parasitoid egg is encapsulated and can be seen as a melanised black area under the cuticle of all subsequent life history stages (Carton et al., 2008). Here we take advantage of melanised areas both on and under the cuticle, as evidence of wounding and successful parasitoid egg encapsulation, respectively.

Wounding therefore exerts two concurrent selection pressures on the immune system: first on a rapid and efficient wound repair, and second on responding to microbial invaders. In this study, we determine the frequency and type of wounds that female and male *D. melanogaster* sustain in the wild. *D. melanogaster* has been widely used in immunity studies (Lemaitre & Hoffmann, 2007) and as an invertebrate model for wound healing in the lab (Belacortu & Paricio, 2011; Dziedziech et al., 2020). Experimental evidence from laboratory studies show that *D. melanogaster* has aggressive territorial interactions with *D. simulans*, where males can end up limping, suggesting damage has been caused (Hoffmann, 1987), and that copulatory wounds are found across many female *Drosophila* species including the melanogaster sub-group (Kamimura, 2007, 2010). Furthermore, wing damage resulted from aggressive behaviour in group-housed *D. melanogaster* males (Davis et al., 2018). There is anecdotal evidence of darkened melanised spots on the cuticle of wild-collected *D. melanogaster* (Chambers et al., 2014). However, there is no systematic study exploring the prevalence of wounding in wild *D. melanogaster*, and thus its potential ecological and evolutionary significance, is unknown.

Marine benthic invertebrates (Lindsay, 2010) and the crustacean, *Gammarus pulex* (Plaistow et al., 2003) can show sex-specific, temporal, and geographic variation in wounding rates, but apart from female copulatory wounding, these factors are relatively unexplored in insects. We therefore systematically collected more than 1,000 male and female *D. melanogaster* across three seasons and three locations and examined them for wounding, and the presence of melanised patches under the abdomen as evidence of parasitoid egg encapsulation. Lastly, as mite mouthparts can potentially penetrate and wound the host cuticle, we collected more than 8,000 flies, including other Drosophilidae species, to assess the frequency with which they are found.

## 2. Materials & Methods

### 2.1 Sampling sites and collection methods

Adult flies were collected from three farms located in and around Berlin: Domäne Dahlem (hereafter denoted as D, 52.45883°N, 13.28901°E), Obsthof Lindicke (hereafter L, 52.38012°N, 12.86828°E) and SL Gartenbau (hereafter G, 52.69889°N, 13.08673°E). The farms are located approximately 30 to 40 kilometres away from each other. Collections were carried out during three time windows between June and October in 2021, i.e., early summer (hereafter ES, 24^th^ June-21^st^ July), late summer (hereafter LS, 24^th^ August-6^th^ September), and autumn (hereafter A, 4^th^-8^th^ October). Due to the weather-dependent nature of successful fly collection, the collections were made over a different number of days during each collection window.

The flies were collected using traps made from 750ml plastic water bottles, which had been cleaned with 70 % ethanol. To allow flies to enter the traps, three holes approximately equidistance from each other, were made in the upper part of the bottle. Per bottle, three 1.5 mL microcentrifuge tubes were prepared so that the bottom portion of each tube was cut off, leaving an opening of approximately 5 mm diameter. One prepared microcentrifuge tube was pushed into each hole in the bottle, with the smaller end of the tube inside the bottle and the larger opening with the cap, located just outside the bottle. The bottom quarter of the bottles were cut off and half-filled with fruit (banana or apples) mixed with dry yeast granules, which had been allowed to ferment overnight, and then the bottle was taped back together, and the traps were placed into the field the following morning. For each collection site 20 to 25 traps were hung in cherry or apple trees (depending upon the season), and next to compost heaps. The traps were placed in the field in the afternoon, and the flies were collected on the following mornings for up to seven days. To collect the flies from the traps, we covered the bottles in black material with a hole the size of the bottle opening, removed the lids of the bottles, and carefully placed a sterile 50 mL falcon tube over the opening, allowing the flies to fly or walk into the tubes. On the occasions where there were few flies in the traps, we additionally used falcon tubes to collect flies directly from the substrate (< 10% of flies were collected in this way). To avoid post-collection damage, we wanted to immediately immobilise the flies after capture. Therefore, for collections we used dry ice or 99 % ethanol, and carefully transported the flies to the laboratory a maximum of three hours after collection. Once in the lab, the flies were stored in 99 % ethanol at -20°C until they were examined for damage. We tested whether our collection and storage methods themselves affected the amount of damage. We found that the dry ice caused antennae and bristle loss we therefore did not include the loss of these body parts as evidence of damage received in the wild. None of the other tested kinds of damage were affected by the storage methods, therefore they were included in the main survey (see Supplementary Information 1).

We captured several Drosophilidae species in our traps, but we aimed to examine only *D. melanogaster* for wounds. Except for *D. simulans* females, both sexes from all other species could be distinguished from *D. melanogaster* based on their morphology. To distinguish between *D. melanogaster* and *D. simulans* females after examining them for wounding, we carried out a diagnostic PCR (see 2.3). The flies that were morphologically identified as neither *D. melanogaster* nor *D. simulans* were kept in ethanol to later examine whether they had mites attached to them. All *D. melanogaster* and *D. simulans* were also examined for the presence of mites.

### 2.2 Examination methods for cuticular damage

We examined a total of 1246 flies for damage: 638 *D. melanogaster* males and 608 females (the latter is a combination of *D. melanogaster* and *D. simulans*). For sample sizes per season and site, see Supplementary Table 1. Flies were examined blind and randomly with respect to season and site. The whole body was carefully examined for melanised spots (Fig. 1), which are an indicator of past damage, or to check for missing body parts (e.g., legs, see Fig.1c). The flies were examined under a Leica M205C stereomicroscope at up to 80 x magnification. Images were captured with a Leica FLEXACAM C1 and Leica Application Suite.

**Figure 1.**
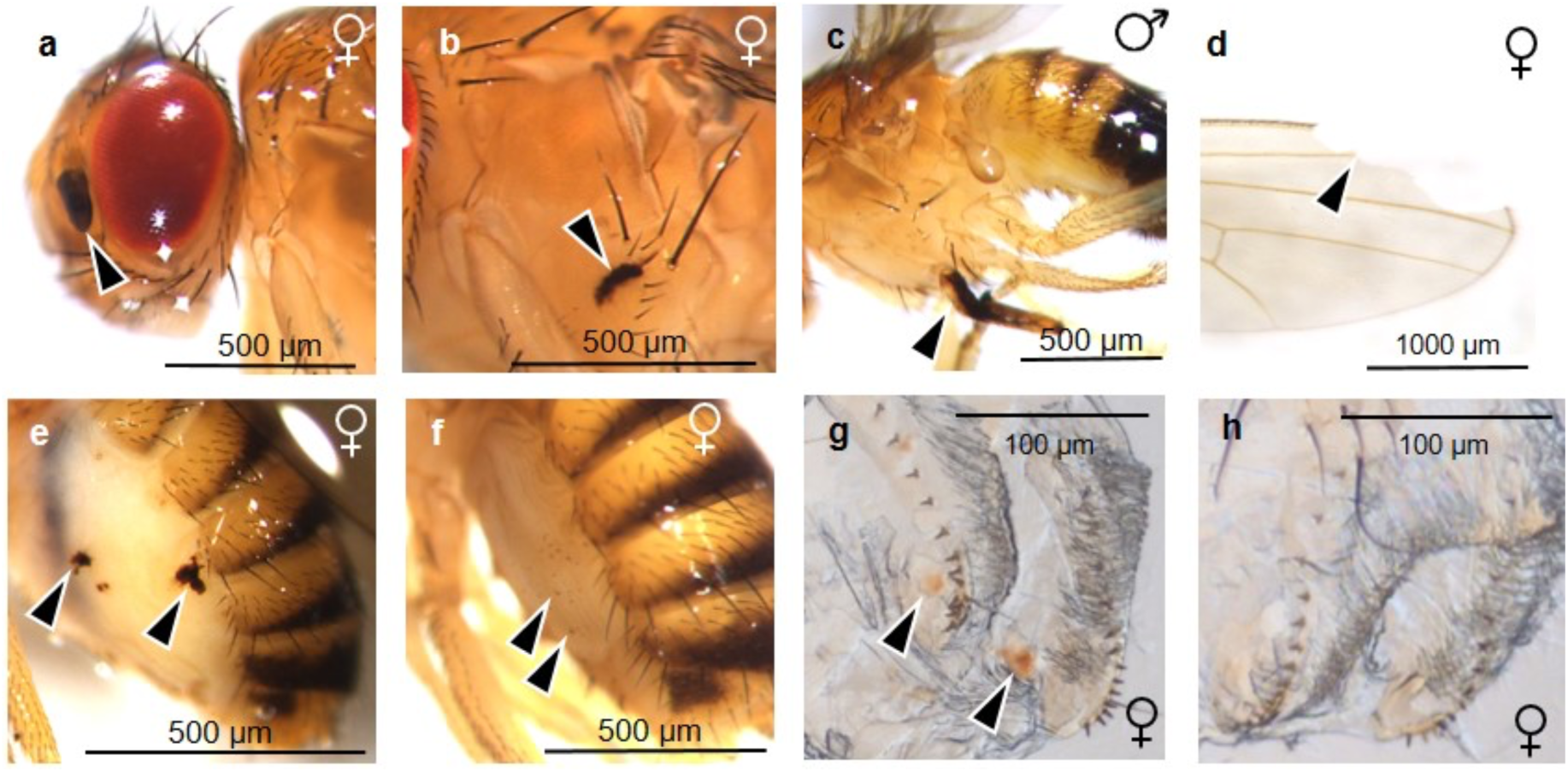
Examples of wounding and damage in *D. melanogaster*. Arrows indicate melanised spots likely resulting from an immune response after cuticular wounding or missing parts of the wings. Wild-collected flies with damage to the (a) head, (b) thorax, (c) leg, (d) wing, (e & f) ventral abdomen. Images from lab-reared flies under controlled mating conditions to illustrate (g) wounds on female vaginal furcal dorsolateral fold (formerly termed lateral folds in Kamimura, 2010, more recent terminology from McQueen et al., 2022) resulting from mating and (h) a virgin female without copulatory wounds. The sex of the fly is indicated on the right of each image. Images (g) and (h) taken by A. Finsterbusch.

#### 2.2.1 Head, leg, thorax, and abdomen wounds and parasitoid eggs

Once the collections were complete, up to 100 samples were examined per day, in a random order with respect to the sex, season, and site, and blind with respect to season and site. The head, thorax, legs, and abdomen were systematically examined, and we used the presence of melanised spots/patches as evidence of wounding (Önfelt Tingvall et al., 2001). For examination, the flies were removed from their tubes and placed on a microscope slide. Throughout the examination process, the surface of the flies were kept wet using a drop of *Drosophila* Ringer’s solution (182mM KCl, 46mM NaCl, 3mM CaCl2.2H2O, and 10mM Tris-HCl, Werner et al., 2000). This method allowed for the easier observation of small melanized patches (see Supplementary Information 1). It is important to note that it was not possible to discriminate between wounds that would likely have penetrated the cuticle and wounds that led to melanisation due to abrasion or scratches on the exterior surface of the cuticle, or melanised spots that may have occurred in the cuticle independently of external damage. We also note that we mostly use the term “wound” but that some literature uses the term “injury” and others “damage” to refer to similar phenomena. The head was examined for melanised patches on the mouthparts, eyes, and antennae (Fig. 1a). Melanised spots on the legs, or missing parts were recorded, and they were combined into the same category for the analyses given their low frequency. For both the thorax and the abdomen we recorded melanised spots on the cuticle (Fig. 1e & f). We also noted large, melanised areas under the cuticle, which is indicative of encapsulated parasitoid eggs. In these cases, the flies were gently squashed between two glass slides and the melanised areas examined under the microscope to differentiate encapsulated eggs from potential internal autoimmune damage (personal communication with Bregje Wertheim).

#### 2.2.2 Wings

We aimed to examine the wings of up to 20 females and 20 males per site and season (total of 334 flies, see Supplementary Table 1); these individuals were randomly chosen from the flies examined in section 2.2.1. The wings were carefully dissected, in a random order and blind with respect to the sex, season, and site. The dissection was performed in *Drosophila* Ringer’s solution by holding the fly with a pair of forceps and removing the wings from the attachment points. The wings were fixed to a microscope slide by using one drop of biological glue, Entellan^TM^ (Merck), and an 18 x 18 mm cover slip was put on top. Photographs were taken under the stereomicroscope mentioned above at a magnification of 64 x with identical light settings for each photograph. The damage on the wings was quantified using similar quantification criteria as in Burkhard et al., (2002), that is (1) notches (small triangular areas at the posterior margin of the wings), (2) tears (where a wing is torn but there is no missing parts) and (3) areas (missing sections of the wing). Considering that haemolymph circulates in the veins (Arnold, 1964; Salcedo & Socha, 2020), we hypothesised that damage to the wing veins might create a possible route for infection. Therefore, each of the above three categories were classified into damage that was, or was not, on the vein, giving a total of six categories (Supplementary Fig. 1). However, due to the limited number of individuals with damage in each category (Supplementary Fig. 2), we instead analysed only whether the wing damage occurred on a vein or not.

#### 2.2.3 Female copulatory wounds

Females were dissected in *Drosophila* Ringer’s solution to assess whether they were internally wounded, which is most likely due to traumatic mating (Kamimura, 2007). The same randomly chosen females used for the wing damage (2.2.2) were used to examine copulatory wounds. For the dissection, the female was held from the tergites on the abdomen with a pair of forceps and a little pressure was applied to the female abdomen to extrude the terminalia. Then a second pair of forceps was used to gently pull the terminalia away from the abdomen. After dissection, the terminalia were placed onto another microscope slide with 5 μl of Ringer’s solution and flattened using an 18 x 18mm cover slip. The vaginal furcal dorsolateral folds were examined under a light microscope (ZEISS Germany Axiophot) at 250 x magnification and photographs were taken, always using identical light settings. Wounds appear as brown spots in mated females, and virgins do not show such wounds (Kamimura, 2007; Fig. 1g & h). We noted the presence or absence of the melanised spots.

### 2.3 D. melanogaster *species identification*

*D. melanogaster* and its sister species, *D. simulans*, commonly share the same habitat and are morphologically only distinguishable from external male genitalia (Sturtevant, 1919). To distinguish female *D. melanogaster* from female *D. simulans,* molecular methods were used. After examining the body for damage, the legs of females were removed using a pair of forceps and used for DNA extraction. We used the single fly genomic DNA extraction method from Gloor & Engles (1992) to extract the DNA. We added a centrifuge step to the protocol after the incubation steps, where the samples were spun down at maximum speed for 1 min and the supernatant was transferred to a new microcentrifuge tube. To distinguish between female *D. melanogaster* and *D. simulans* a PCR was performed using the *Slif* primers as described by Faria & Sucena (2017) except that we used KAPA HiFi HotStart ReadyMix PCR kit (Roche). The gel images were examined to determine the species using the lengths of the amplified fragments: 939 bp for *D. melanogaster* and 1058 bp for *D. simulans*.

### 2.4 Mites

In total 8019 (7612 *D. melanogaster*/*D. simulans*, 407 other Drosophilidae species) flies were examined for the presence of mites. We noted the number of attached mites, where they were attached to the fly body, as well as the season and collection site. However due to unequal sampling across seasons/sites, we did not include those variables in our statistical analyses. The species of the flies with mites attached to them, and the mites, were identified by using molecular methods or morphologically by Darren Obbard by using photographs. Where possible, we determined the sex of the flies.

#### 2.4.1 Molecular identification of fly and mite species and host wound response

Mites were carefully removed from the fly body under the stereomicroscope using a pair of forceps. We noted whether melanised spots were present where the mouthparts were attached to the mite and took photographs under the stereomicroscope. DNA extraction and PCRs were carried out following the methods as described in Perez-Leanos et al. (2017) with a slight modification to the homogenisation step. Briefly, the DNeasyTM (QIAGEN) DNA extraction kit was used to perform the DNA extraction from the flies and the mites. After being separated, each fly and mite were put into separate 1.5 mL microcentrifuge tubes which contained 180 µl ATL buffer. The mites were pooled if more than one morphologically identical mite was found on one fly. To homogenise the flies and the mites, we used two tungsten carbide beads (3 mm, QIAGEN) and the homogenisation was done in a Retsch Mill (MM300) at a frequency of 30 Hz for 5 min. The rest of the DNA extraction was performed according to the manufacturer’s protocol. The same PCR conditions and primers were used as in Perez-Leanos et al. (2017) and sequencing was performed at Eurofins Genomics, Germany. The sequences were aligned using BLASTN in the NCBI genome browser. All sequencing data will be deposited in GenBank.

#### 2.4.2 µCT and SEM

To examine whether the mouthparts of the mites were piercing the fly cuticle, and to examine encapsulated parasitoid eggs we performed X-ray microtomography (µCT). For this we fixed three flies with mites and one with an encapsulated parasitoid in 4 % formaldehyde in 80 % ethanol for two to three days at room temperature. After two times one hour washing with 80 % ethanol the samples were transferred to 100 % denaturated ethanol for two to three days and stored at room temperature, which increases the latter contrast in the µCT. Next, samples were further contrasted in a 1 % methanolic iodine solution for one day at room temperature. After that the samples were rinsed three times with 100 % denatured ethanol and three times with 100 % pure ethanol for one hour each step. Finally, samples were transferred to the critical point dryer CPD 300 (Leica, Nußloch, Germany). The dried samples were glued with UV curing glue to a holder for inserting them into the µCT SkyScan1272 (Bruker, Billerica MA, US). CT scans were acquired with x-ray source running at 60 kV and 130 uA. Pixel scaling is 1.5 µm in xyz with a rotation step of 0.2° and an image size of 2059 x 1640. Acquisition was controlled with the setup software and reconstruction of the scans with ring artefact correction was done within NRecon (Bruker, Billerica MA, US). Visualisation of the final scan was done using Dragonfly (Object Research Systems (ORS), Montréal, Canada). In addition, we acquired images with the scanning electron microscope (SEM) to get a closer look of the attachment sites of the mites, where possible. Therefore, the critical point dried samples were sputtered with gold after the µCT scan and were visualised using a LEO 1450 VP (Zeiss, Oberkochen, Germany) running with 8 kV.

### 2.5 Replication Statement

We wish to understand how sex, season, and site affect wounding and parasitoid encapsulation in *D. melanogaster*, and how season and site affect wounding and parasitoid encapsulation in female *D. simulans* (Table 1). The sample sizes for each combination of factors are in Supplementary Table 1. We also wish to understand whether sex, season and site affect the presence of mites in Drosophilidae (Table 1). See also Supplementary Table 2.

**Table 1.**
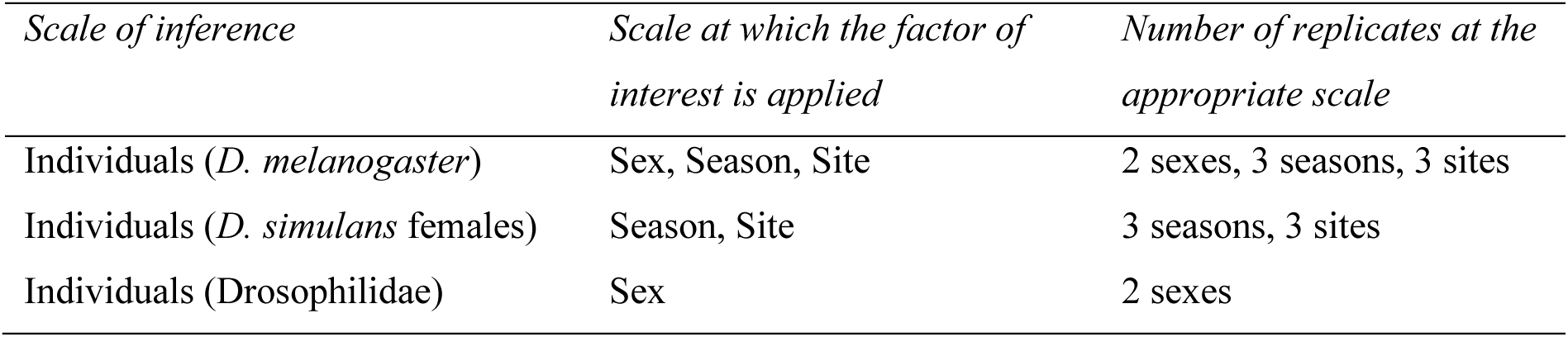
Replication statement for the questions addressed in this study.

| *Scale of inference* | *Scale at which the factor of interest is applied* | *Number of replicates at the appropriate scale* |
| --- | --- | --- |
| Individuals (*D. melanogaster*) | Sex, Season, Site | 2 sexes, 3 seasons, 3 sites |
| Individuals (*D. simulans* females) | Season, Site | 3 seasons, 3 sites |
| Individuals (Drosophilidae) | Sex | 2 sexes |

### 2.6 Statistical analyses

Analyses were performed using R version 4.2.2 (R Core Team, 2023) in RStudio version 2022.07.2. For the generalised linear models, we tested main effects and two-way interactions. Unless stated otherwise, the “DHARMa” (Hartig, 2022) package was used for residuals diagnostic of the statistical models, analysis of variance tables were produced using “car” (Fox & Weisberg 2019) using type III ANOVAs in the presence and type II ANOVAs in the absence of interactions, and post-hoc tests were carried out with “emmeans” (Lenth, 2023) using the Tukey adjustment. To conduct generalised linear mixed models, “glmmTMB” package (Brooks et al., 2017) was used. To visualise the data “ggpubr” (Kassambara, 2023), “ggplot2” (Wickham, 2016) and “viridis” (Garnier et al., 2023) were used. Transformation and manipulation of the data were carried out with “tidyverse” (Wickham et al., 2019). All the statistical tests mentioned below were performed for both *D. melanogaster* and *D. simulans*, and details of the statistical analysis for *D. simulans* can be found in Supplementary Information 2.

#### 2.6.1 Head, legs, thorax, and abdomen damage

We first tested whether there is variation in the number of body parts that are damaged per fly. To do this we performed a Chi-square test using the number of individuals with none, one, two, three or four body parts damaged. The four body parts considered were the head, legs, thorax, and abdomen. Post-hoc multiple comparisons were not performed because of the low numbers of flies with three and four injured body parts, i.e., five and three fly respectively.

To test whether the body parts differed in their likelihood of being damaged, we performed a Chi-square test with the number of individuals with and without damage to each of the head, leg, thorax, and abdomen. Post-hoc multiple comparisons were performed with “fifer (Fife, 2014)”, using the Bonferroni adjustment for multiple comparisons.

Given that relatively few flies had damage to more than one body part (Fig. 2a), we produced a new response variable called “combined damage HTLA” (head, legs, thorax and abdomen), that collapsed damage into one variable: a “0” means that a fly had no damage, and a “1” means that the fly had damage to one or more body parts. We tested whether combined damage HTLA was affected by sex, season or site and their two-way interactions, using a generalized linear model (glm) with binomial distribution:

**Figure 2.**
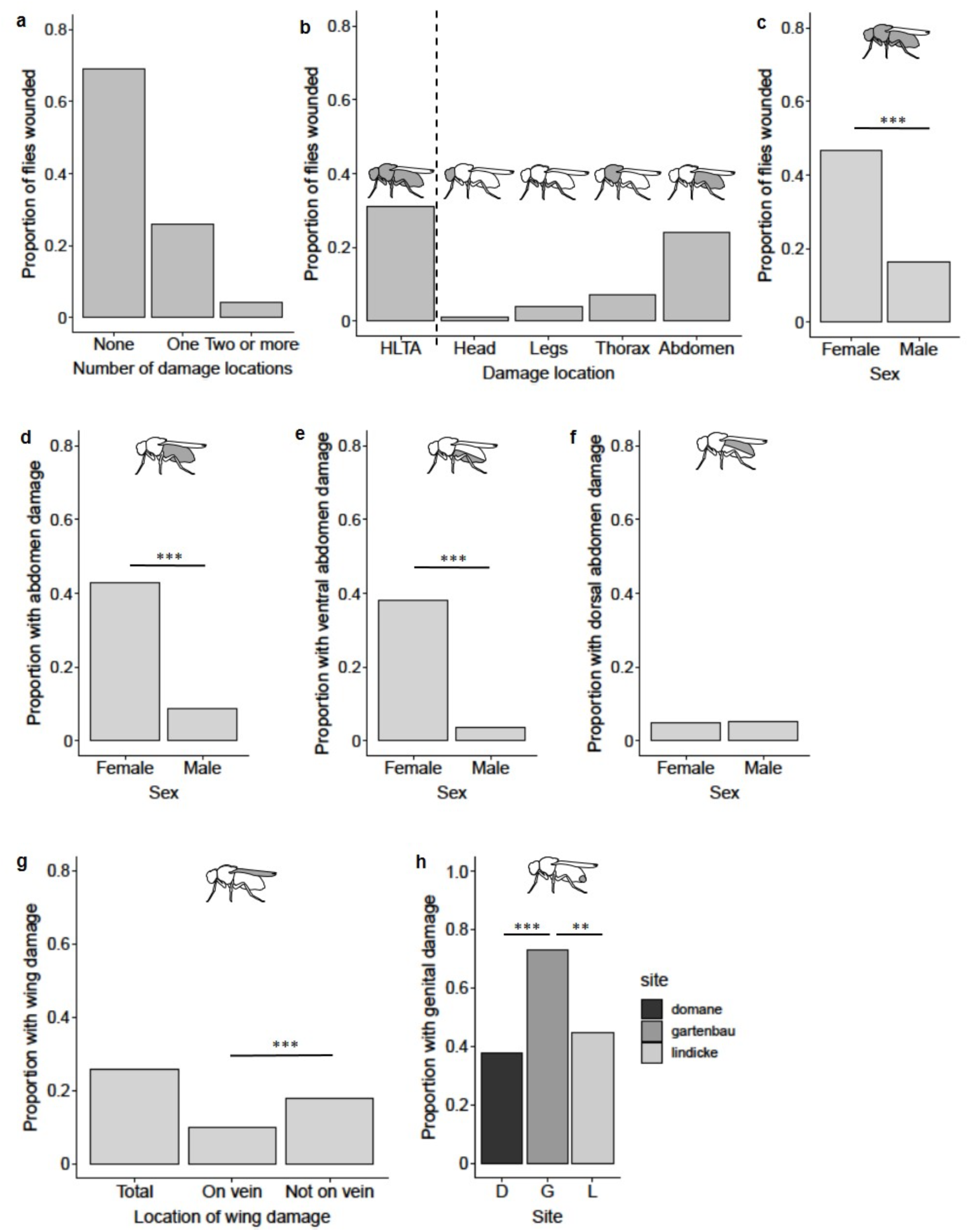
Frequency of wounding in wild-collected *D. melanogaster*. (a) The proportion of flies with none, one or more than one type of damage, (b) the proportion of flies with wounds to the external cuticle (HTLA: head, legs, thorax and abdomen), (c) the proportion of females and males showing HTLA wounding, (d-f) the proportion of the flies with total, ventral and dorsal abdominal melanised spots by sex (for panels a-f, n = 1174), (g) the proportion of flies with wing damage (total) and damage that is either to a vein or not to a vein (n = 338), and (h) the proportion of females showing copulatory wounding according to collection site (n = 163). Stars indicate statistically significant groups where: ** = p <0.001 and *** = p <0.0001.

Model 1: combined damage HTLA ∼ sex × season + sex × site + season × site

We then examined individual body parts separately, and tested whether sex, season or site affected the number of flies with thorax wounds, abdomen wounds, or wounds to the ventral or dorsal abdomen. We used the same model as Model 1, but the binary response variables were thorax damage, total abdomen damage, or ventral or dorsal abdomen damage. Furthermore, we investigated whether the frequency of ventral abdomen damage differed from dorsal abdomen damage by using a Chi-square test. The head and legs were not tested because of the low proportion of flies with damage to these body parts (Fig. 2b).

#### 2.6.2 Wing Damage

To test if there was a difference in the proportion of flies with wing damage that did or did not include wing veins, we performed a Chi-square test. We tested the effect of sex, season and site on damage that affected the veins, and in a separate model, on damage that did not affect the veins. Once again, using Model 1 but replacing the response variable.

#### 2.6.3 Female genital damage

We asked whether the season, site, or an interaction between these two factors, affected the number of females with genital damage, by using Model 1 but replacing the response variable. Female and male damage to the abdomen was mostly observed on the ventral abdomen, and females had this damage more frequently than males (see results). As a result, we hypothesised that the melanised spots on the female abdomen could be obtained during copulation. Therefore, we tested whether there was a relationship between the presence of genital damage and the presence of melanised spots on the abdomen. For this we used a binomial glm with genital damage as the response variable and abdominal melanized spots as a covariate.

#### 2.6.4 Encapsulated melanised parasitoid eggs

To test whether the proportion of flies with encapsulated melanised parasitoid eggs, was affected by sex, season, or site, we used binomial glm with presence or absence of abdominal parasitoid as the response variable and sex, season, and site as a factor in two-way interactions as in Model 1.

#### 2.6.5 Mites

Using Chi-square tests, we asked whether there is variation in the number of mites attached per fly. Three samples were excluded from this analysis due to unknown attachment sites. We also tested whether mites are attached to some fly body parts more frequently than others. To do this we again used a Chi-square test. After the removal of the mites from the fly body, we sometimes observed melanised spots where the mouthparts had been. Therefore, by using a one-sample proportion test, we tested whether there is a difference in the frequency of observed melanised spots. To investigate the susceptibility of females and males to mite infestation, we employed generalised mixed models (glmmTMB) with a betabinomial distribution. The response variable, representing the presence and absence of mites, was combined into a single object using cbind. In the analysis, sex, season and site were considered fixed factors. The data used for this test consisted of a table detailing the number of mite-infested/not infested females and males among the total collected individuals in each season and site.

Model 2: cbind (with mites, without mites) ∼ sex + season + site

## 3. Results

In total 1246 flies were examined for any type of wound or damage, of which 638 were morphologically identified as male *D. melanogaster*. Diagnostic PCRs allowed us to identify 536 female *D. melanogaster* and the remaining 71 females were *D. simulans*. The latter were analysed separately from *D. melanogaster* (see Supplementary Information 2 for *D. simulans* results). We examined six areas of the body for damage (head, leg, thorax, abdomen, wing, and female genitalia; Fig. 1).

### 3.1 Head, leg, thorax, and abdomen damage

There was significant variation in the number of wounded body parts per fly (Chi square = 769.81, df = 2, p < 0.0001, n = 1174). Flies most frequently showed wounds on one body part, with only a small percentage of the flies having two or more wounded body parts (Fig. 2a). The maximum number of wounded parts was four, which was found in only three flies. Thirty one percent of individuals showed at least one type of wound on the external cuticle (Fig. 2b). The head, legs, thorax, and abdomen differed significantly in the frequency with which they were wounded (Chi square = 437.09, df = 3, p < 0.0001, n = 1174), with the abdomen most frequently showing damage (Fig. 2b).

When we combined damage across the head, thorax, legs, and abdomen into one response variable, we found that females were more frequently injured than males (Supplementary Table 3, Fig. 2c). There was also a significant interaction between site and season (Supplementary Table 3), and multiple comparisons showed that this was due to late summer differences, where flies from the D site were less frequently damaged compared to those from G and L.

Season and collection site significantly affected the proportion of flies with thorax damage (Supplementary Table 4). Post-hoc multiple comparisons showed that the early summer flies less frequently had damage to their thoraces compared to late summer (z = -2.82, p = 0.0134). Females were significantly more likely to have damage on their abdomens compared to males (Supplementary Table 5; Fig. 2d). There was also a significant interaction between season and site; after multiple comparisons, late summer flies from site D had significantly more damage on their abdomen compared to those from sites G (z = -3.154, p = 0.0131) and L (z = -3.690, p = 0.0069). The ventral abdomen was more frequently wounded compared to the dorsal abdomen (Chi square = 96.50, df = 1, p < 0.0001, n = 1174; Fig. 2e & f). Furthermore, females were more likely to have damage to their ventral abdomen compared to males (Supplementary Table 5) and there was a significant interaction between season and site. After multiple comparisons, it was revealed that early summer flies from site D had significantly more damage than late summer flies from site D (z = 3.116, p = 0.0480), and late summer flies from site D had more damage than late summer flies from sites G (z = -3.321, p = 0.0252) and L (z = -4.004, p = 0.0020). However, there was no significant effect of any factors on the damage to the dorsal abdomen (Supplementary Table 5).

### 3.2 Wing damage

In total 338 flies (165 females, 173 males) were examined for wing damage. Damage was less frequently found to affect a vein than part of the wing not containing a vein (Chi-square = 12.81, df = 1, p = 0.0003; Fig. 2g). When we only considered wing damage that affected veins, we found a significant difference between seasons and sites (Supplementary Table 6) but after multiple comparisons, these differences no longer remained significant. Season was the only factor to affect the wing damage not on the veins (Supplementary 6) and multiple comparisons showed that this was due to late summer differences compared to the autumn.

### 3.3 Copulatory wounds

In total 178 females were examined for copulatory wounds, and of those 163 were molecularly identified as *D. melanogaster* and 15 as *D. simulans*. Fifty two percent of *D. melanogaster* showed copulatory wounds, and the proportion varied significantly with collection site (Supplementary Table 7, Fig. 2h). We hypothesised that abdominal damage found on females might be related to mating, and there was a marginally non-significant positive relationship (LR Chisq = 3.74, p = 0.0527) between the presence of copulatory and abdominal wounds. However, when focusing only on damage to the ventral abdomen, no significant correlation was detected.

### 3.4 Encapsulated melanised parasitoid eggs

Melanised parasitoids were predominantly observed in the abdomen, with one found in the head and another in the thorax (Fig. 3a-c). A melanised parasitoid egg was visible through the cuticle of 9.7 % (114 out of 1174) flies. Season and site together affected the proportion of flies with parasitoids (Supplementary Table 8; Fig. 3d), which was driven by differences between the early summer L site and late summer G site (z = -3. 22, p = 0.0346). There was also a significant interaction between sex and site (Supplementary Table 8) although after multiple comparisons none of the groupings differed from each other. The number of encapsulated parasitoid eggs in a fly differed significantly (Chi square = 200.32, df = 4, p < 0.0001, n = 1174, Fig. 3e). In 70 % of the flies, a single encapsulated parasitoid egg was observed and the rest of the flies containing two or more encapsulated parasitoid eggs.

**Figure 3.**
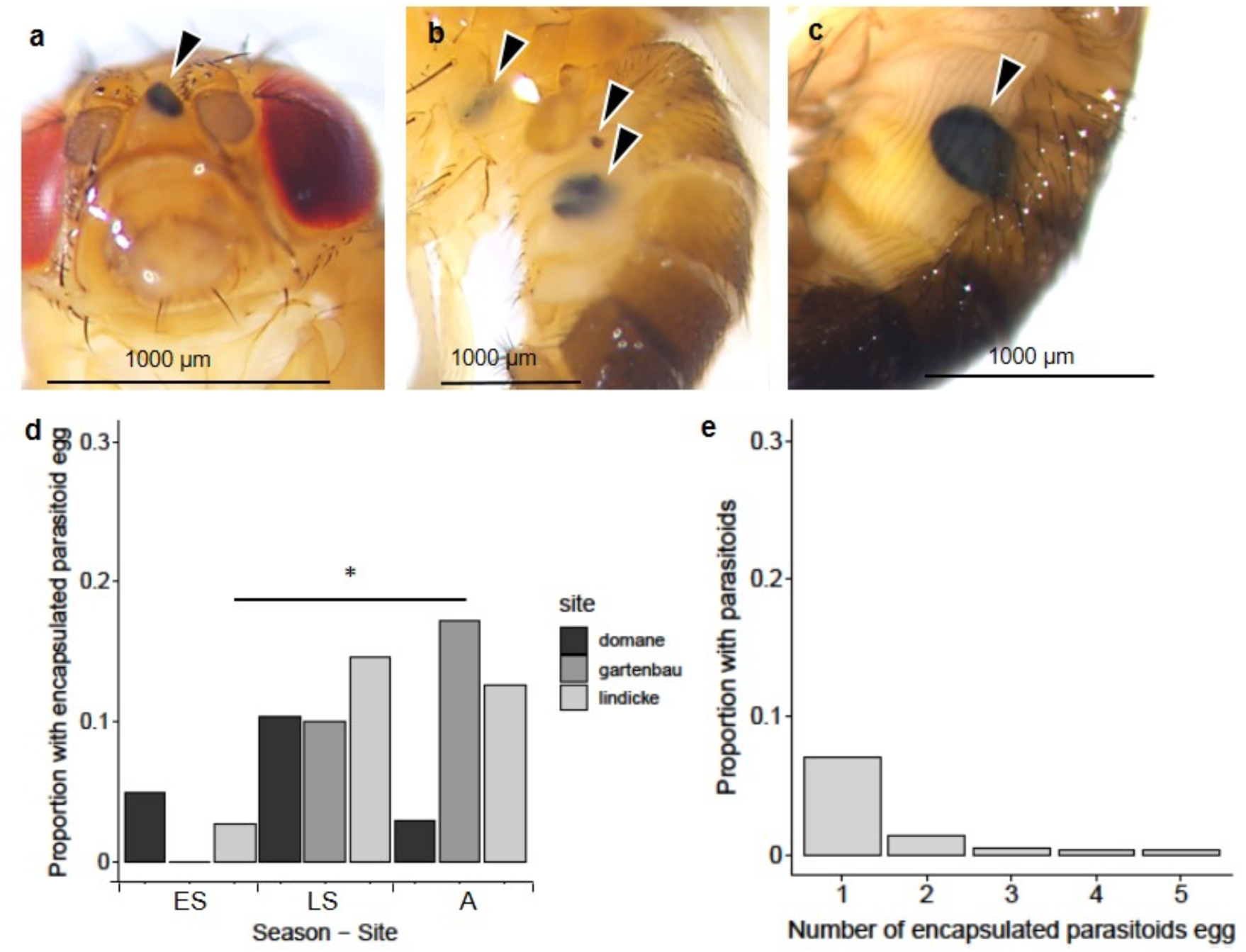
Melanised encapsulated parasitoid egg in wild-collected *D. melanogaster*. Images of melanised encapsulated parasitoid egg in (a) the head, (b) the thorax and abdomen, and (c) the abdomen. (d) the proportion of flies with parasitoids by season and site and (e) the proportion of flies with one or more encapsulated parasitoids (for panels d & e, n = 1174). ES: early summer, LS: late summer and A: autumn. Stars indicate statistical significance, where * = p < 0.01.

### 3.5 Mites

In 0.7 % (56 of 8019) of the collected flies, one or more mites were attached to the fly body (Fig. 4a). Mites were found on seven Drosophilidae species: *Drosophila busckii, Drosophila hydei, D. melanogaster*, *D. simulans*, and *Drosophila subobscura* which were identified via sequencing, and *Drosophila funebris and Scaptomyza pallida*, which were identified phenotypically (Supplementary Table 2). We identified 18 out of 56 mites via sequencing; two mites were identified to the genus level: *Macrocheles sp*. (Fig. 4b) and *Pergamasus sp* (Fig. 4c), and one mite was identified to the species level: *Archidispus insolitus*. The remaining mites were not identified molecularly due to unsuccessful DNA extraction, or the sequences had the highest similarity to an organism other than a mite. Among the identified mites, 14 of the flies were found to be parasitised with *Macrocheles sp.* while *Pergamasus sp.* was found on three *D. melanogaster* and one *D. subobscura*; *A. insolitus* was found on one *D. subobscura* (Supplementary Table 2).

**Figure 4.**
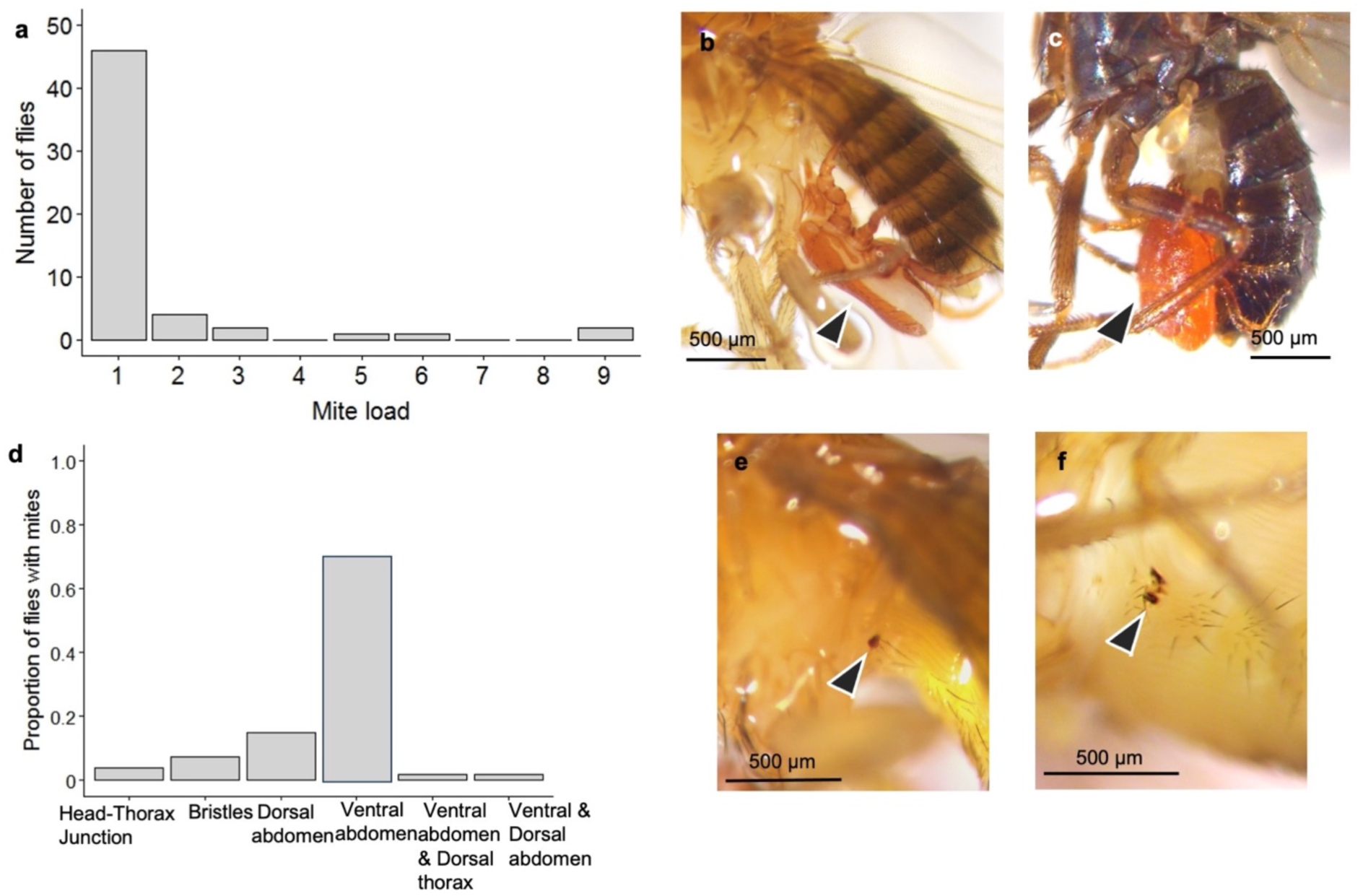
Mites found on wild-collected Drosophilidae. (a) mite load across flies, (b) *Macrocheles sp.* attached to a female *D. melanogaster*, (c) A *Pergamasus sp*. attached to a *D. subobscura*, (d) the numbers of mites attached to different body parts. The bristles were on the head and the thorax. One fly had mites attached to both the ventral abdomen and dorsal thorax, and another fly was found with a mite attached to both the ventral and dorsal abdomen. (e & f) the melanised area visible on the fly body after removing the mite.

There was significant variation in the number of mites that were attached to a fly (Chi square = 173.5, df = 5, p < 0.0001, N = 56; Fig. 4a), and their attachment sites (Chi-square = 111.72, df =5, p < 0.0001, N = 53; Fig. 4d), together indicating that most flies had one mite attached to the ventral abdomen. After removing the attached mites, 74.4 % (32 out of 43 flies) more flies had one or more melanised patches compared to no melanised patches (Chi square = 9.30, df = 1, p < 0.0023, N = 43; Fig. 4e & f). Sex, season, and site had no significant effect on the probability of having an attached mite (Supplementary Table 9).

### 3.6 µCT and SEM

The µCT of three flies indicates that the mites attached to flies in different ways (Fig. 5). The mites in Fig. 5a appear to penetrate the fly cuticle, in combination with the presence of a mite- or fly-derived substance (Fig. 5a””), which is indicated by a more x-ray dense and therefore brighter contact site between mite and fly (Fig. 5a”), whereas the larger mite in Fig. 5b seems to grasp an abdominal cuticular fold with their chelicerae. The other mites appear to cement themselves to the abdomen without cuticular penetration, again visible by the bright contact site in the µCT visualisation (Fig. 5c). In the case of the encapsulated endoparasitic wasp eggs, the µCT shows the remains of the capsule inside the fly body (Fig. 5d); it appears to consist of two parts, one is an outer dome shaped hull and the second structure is a more diffuse matter inside of the hull.

**Figure 5.**
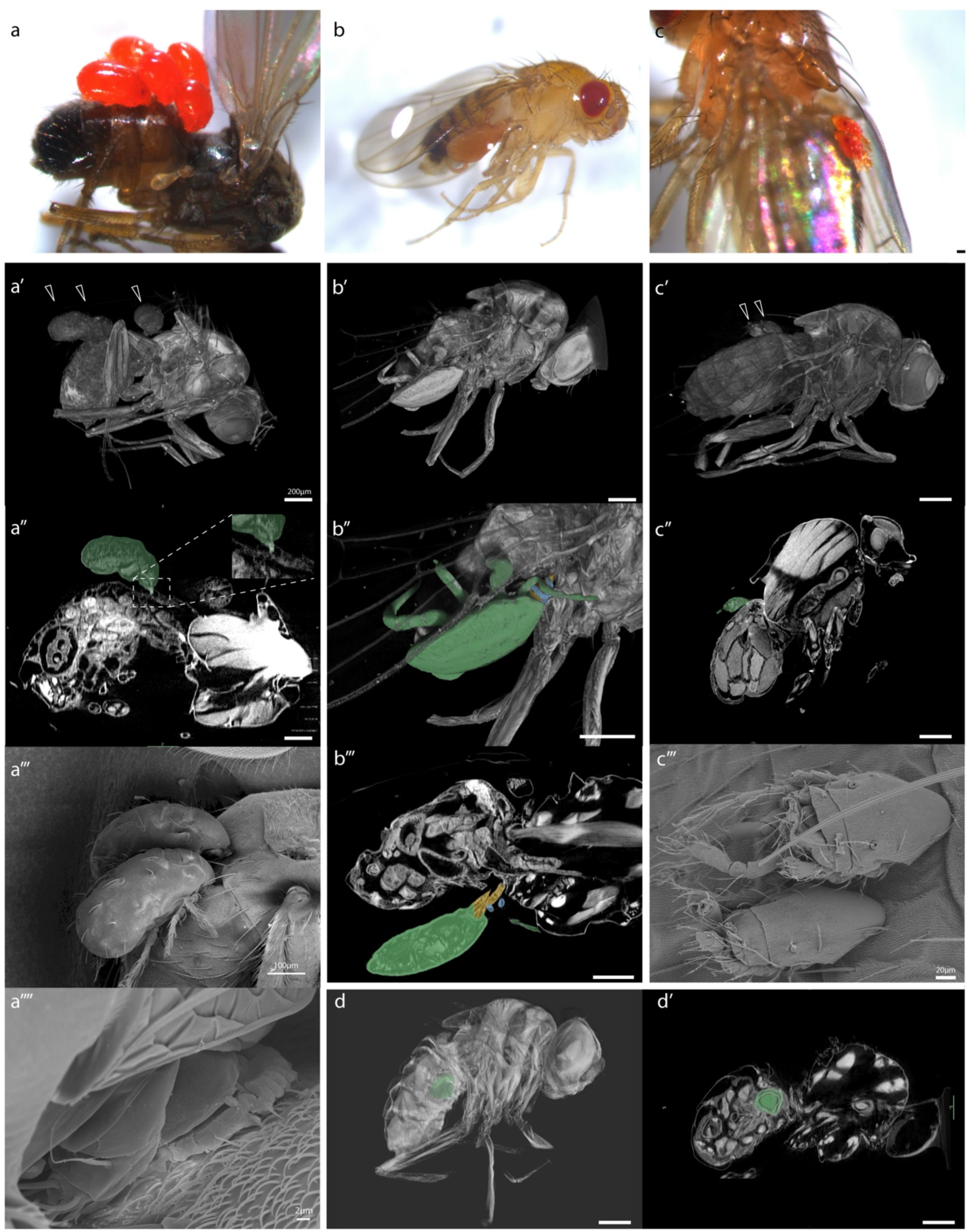
µCT and SEM of mites attached to Drosophilidae and an encapsulated parasitoid. **(a)** *S. pallida* with three mites (arrowheads, green) whose chelicerae slightly penetrate the dorsal abdominal cuticle and where a mite- or fly-derived substance is visible at the contact point. (b) *D. melanogaster* or *D. simulans* female with a single large mite (green) attached to the ventral side of the abdomen, grasping a cuticular fold with the chelicerae (yellow) and pedipalps (blue). (c) Two smaller mites (arrowheads, green) cemented to the dorsal side of a *D. melanogaster* or *D. simulans* female abdomen. (d) Remnant of an encapsulated endoparasitic wasp inside a *D. melanogaster* male, which is visible in two parts (hull – light green, inside – dark green). Scale bars indicate 200µm if not otherwise indicated.

## 4. Discussion

The immune defences of *D. melangoster* are remarkably well-studied in the lab, yet our knowledge from the field is lacking. Wounds elicit immune responses, and they are potential entry points for pathogens, it is therefore important to understand their frequency in the wild. We find that wounding is prevalent in the wild *D. melanogaster* populations that we studied, with the abdomen being the most frequently wounded body part, particularly in females. When considering interactions with other species that can damage the host, encapsulated parasitoid eggs were found in just under 10 % of individuals. Furthermore, across seven Drosophilidae species, just under one percent carried mites, and we found that most of these mites also wounded their host. Importantly, we note that we may have under-sampled wounded or parasitised individuals because they could have a lower survival, or they might be less likely to have made it into our traps, compared to non-wounded and non-parasitised individuals.

Approximately 31 % of *D. melanogaster* and 36 % of *D. simulans* females had at least one wounded body part, which falls within estimates from studies on other insect species: for example damage or wounding was observed in less than 10 % of male horned beetles (*Allomyrina dichotoma*; Siva-Jothy, 1987) and 17 % of male giant rhinoceros beetles (*Trypoxylus dichotomus*; McCullough, 2014), and wounding can be inferred in up to almost 100 % of two Eurasian Bluets damselflies (*Coenagrion puella* and *Coenagrion hastulatum*), as a result of ectoparasitic mite prevalence (Rolff, 2000). We note that non-sterile abrasion of *D. melanogaster* larval cuticle resulted in melanised marks and activated immune gene expression in the epidermis (Önfelt Tingvall et al., 2001), so even if our study did not distinguish between penetrant and more superficial wounds, the latter may still be immunologically relevant if adults also show such a response. We found a sex-specific difference in susceptibility to cuticular injury, which might be attributed to differences in behaviour or physiology. For example, if females have a longer lifespan in the wild, they may encounter more opportunities to be wounded, and secondly the larger body size of females might give them a higher likelihood of being wounded. The sex difference in wounding appears to be driven by damage to the ventral abdomen. More damage to the ventral abdomen might be explained by the differences in rigidity between the ventral and dorsal abdomen. We hypothesised that female abdominal damage could result from mating, although there was only a weak non-significant positive relationship between the presence of ventral abdominal wounding and copulatory wounding. Another possible explanation for the presence of melanised spots on the ventral abdomen is that this is the dominant mite attachment site, a finding that aligns with other *Drosophila* studies (Polak, 1994; Perez-Leanos et al., 2017; Michalska et al., 2023). Melanised wounds were observed in three-quarters of the flies that had the mites removed, and wounds with a similar appearance were found in our wounding survey flies. However, the frequency with which we found mites attached to flies is considerably lower than the frequency of abdominal wounding and given that there was no effect of sex on the presence of the mites, this may not explain the sex difference in ventral abdominal wounds.

Wing damage was found to less frequently affect an area containing a vein compared to areas without veins, indicating that certain areas of the wing are more prone to damage than others. Given results from a lab study on *D. melanogaster* (Davis et al., 2018) we had hypothesised that males would incur wing damage, and we indeed found it in both males and females. Collection site and season affected wing damage on veins and thorax damage, suggesting that environmental factors may play a role in determining the incidence of injuries. It has been shown that wing injuries in yellow dung flies, *Scathophaga stercoraria*, vary between seasons and relate to the increase in male activity and/or longer female pre-productive periods (Burkhard et al., 2002). These results highlight the importance of considering environmental factors when studying the distribution and prevalence of wounding in natural populations. However, we could not control for age in our experiment, so if older flies are more likely to be wounded than younger flies (e.g., wing damage increased with age in lab *D. melanogaster*; Davis et al., 2018), an alternative explanation is that we may have trapped flies of different ages across seasons and sites. Given that in *D. melanogaster* successful mating results in a copulatory wound (Kamimura 2007 and unpublished data) and that the prevalence of copulatory wounding in the wild varied from ∼35 to ∼75 % across collection sites, it suggests that the numbers of virgin and mated females that we collected differed across sites. In a lab study, 80 % of female *D. melanogaster* were found to reach reproductive maturity at four days post eclosion (Pitnick et al., 1995), which if this is also the case in our population it might suggest that we collected relatively recently emerged flies at the two sites with a lower frequency of copulatory wounding, or that mating opportunities were low. In the lab, bacteria can be transferred from male *D. melanogaster* to the female during mating, resulting in female death (Miest & Bloch-Qazi, 2008). Therefore, in the wild the consequences of genital wounds might have a serious impact on survival. Wounds to the head, thorax and legs were less frequently observed compared to the abdomen and wings. We only collected living flies, which must therefore have survived any injuries that they had sustained. One possible hypothesis to explain variation in the body parts likely to be wounded is that some wounds are more severe and result in higher mortality than others, for example, the proportion of the flies that survived thorax wounds was lower than the flies surviving abdominal wounds (Chambers et al., 2014).

Approximately 42 hymenopteran species have been reported as endoparasitoids of *Drosophila* species (Carton et al., 1986). In addition to wounds, we found that just under ten percent of flies contained one or more melanised parasitoid eggs. Similar to estimates for the frequency of wounding, the frequency of the presence of encapsulated parasitoids in the wild is quite variable. For example, between 39 and 85 % for *Leptopilina heterotoma* and *L. boulardi* in Tunisian *D. melanogaster* and *D. simulans* (Rouault, 1979, referenced in Carton et al., 1986) and between 0 and 50 % of Dutch *Drosophila* species had encapsulated parasitoids (de Haan et al., 1987). There are also numerous estimates on the proportion of encapsulating hosts in the lab, where unlike in the field, it is possible to know the number of cases where encapsulation was nosuccessful, i.e., when the fly died because of the parasitoid. The proportion of encapsulating flies is also highly variable in these lab studies, and includes a population sampled from Berlin where ∼ 60-70 % of *D. melanogaster* successfully encapsulated parasitoid eggs (e.g., Gerritsma et al., 2013). We also observed variation in the prevalence of encapsulated parasitoids between seasons and locations, which aligns with previous work showing that the encapsulation rate can vary among fly populations based on geographical location or seasonal changes, influenced by factors such as host-parasitoid interactions, abiotic and biotic factors (Fleury et al., 2004; Gerritsma et al., 2013).

Lastly, we examined the potential for mites to cause wounds. It has been shown in *Drosophila nigrospracula* that *Macrocheles subbadius* can pierce the fly cuticle and feed on their haemolymph (Polak, 1996), and act as parasite vectors, for instance, *Varroa destructor* can transmit *Deformed wing virus* (DWV) and *Acute bee paralysis virus* (ABPV) in wild honey bee colonies (*Apis mellifera*) and the endosymbiont *Spiroplasma* can be transmitted horizontally with *Macrocheles* species in *Drosophila* (Horn et al., 2020; Jaenike et al., 2007; Murray et al., 2020; Osaka et al., 2013). Fifteen Drosophilid species including some of those that we collected, i.e., *D. melanogaster*, *D. simulans*, *D. busckii* and *D. hydei* have been found infected with ectoparasitic mites in their natural habitat (Jaenike et al., 2007; Polak, 1996, Perez-Leanos et al., 2017). In our study, we identified three Drosophilidae species, *D. subobscura*, *D. funebris and S. pallida* which have not previously been reported to be parasitied by mites. We also identified two mite species, *Archidispus insolitus* and *Pergamasus sp.,* which have not previously been reported on *D. melanogaster*, in addition to the previously reported *Macrocheles sp* (Polak, 1996, Perez-Lanos et al., 2017). We found mites attached to 0.7 % of our collected Drosophilidae, a proportion that is remarkably consistent with the overall proportion of mites found on flies collected using similar methods to our study in Mexico and southern California (Perez-Leanos et al., 2017). It is important to note that many mite species detach themselves from their insect hosts when they become fully engorged, therefore we might have underestimated mite prevalence.

Furthermore, if a mite load was particularly heavy, or if indeed a fly was significantly wounded, it may not have been collected by our trapping method. The µCT images suggest that at least one mite species penetrates the fly cuticle with its mouthparts, but in the other cases there was no obvious penetration. This could be due to there being only shallow penetration between the mite and host that cannot be captured with the methods we used, or because the species that we examined are phoretic.

Overall, the results of this study provide insights into the prevalence and distribution of wounding in natural *D. melanogaster* populations. We show that wounds are frequent in the wild. Wounding, as a common stressor, can lead to physiological changes that influence an individual’s energy allocation, reproductive success, and overall survival; it requires not only a prompt and effective wound healing process but also necessitates an efficient immune response to counter potential infections that may arise from the wounds. Additionally, the presence of ectoparasites like mites poses further challenges, as they can exacerbate the effects of wounds and contribute to disease spread. Therefore, further research is necessary to gain a comprehensive understanding of the impact of mites, other parasites, predators, competitors and conspecifics on *Drosophila* populations. Given the prevalence of wounding in our systematic study, and reports from the literature on wounding in insects and other animals, it suggests that wound repair is almost certainly an important driver of the evolution of immune systems.

## Acknowledgements

The authors thank Darren Obbard and Bregje Wertheim for advice on fly species identification and parasitoids respectively, Mathias Franz for statistical advice, and Diana Aldana Alvarez and Lukas Maas for technical assistance, and Mrs. and Mr. Lindicke, Jens Scheidereit and Annika Siegel for allowing us to sample at the farms, and Anika Finsterbusch for images used in Figure 1. SAOA was supported by a Heisenberg Fellowship (AR 872/4-1 and AR 872/7-1) from the Deutsche Forschungsgemeinschaft (DFG; https://www.dfg.de/en/index.jsp). BSS was supported by the DFG through project AR 872/5-1, awarded to SAOA as part of the Research Unit FOR 5026 “InsectInfect”.

## Author contributions

SAOA conceived the overall idea with input from BSS and JR; BSS, VG, MK and SAOA designed methodology; BSS collected all data except for µCT images, which were produced by VG; BSS, VG and SAOA analysed the data; BSS, VG, MK, JR and SAOA interpreted the data; BSS and SAOA led the writing of the manuscript. All authors contributed critically to the drafts and gave final approval for publication. Our study brings together authors from three countries, including scientists based in the country where the study was carried out.

## Data availability statement

Upon publication, the datasets generated in this study will be made available on Refubium (https://refubium.fu-berlin.de/), the institutional repository of the Freie Universität Berlin.

