## Supplementary files for "How frequently are insects wounded in the wild? A case study using *Drosophila melanogaster*"

### Supplementary Information 1

**Supplementary collection method experiments**

We carried out three experiments to test if the storage methods directly after collection in the field and before placing in ethanol for longer-term storage, caused damage to the fly. For all experiments, we used five- to six-day old *D. melanogaster* adults collected from a population cage that contained the offspring (minimum F8) of wild-collected flies. The cage is maintained at a population size of around 5,000 individuals.

***Experiment 1****: Testing the effect of storage methods on bristle and antenna loss and damage to legs and wings*

In the first experiment, we asked whether the post-collection storage methods that we used resulted in the loss of head bristles, thorax bristles, or antennae, whether they or caused damage to legs or wings. We tested dry ice, ethanol, and dry ice plus ethanol, as all these methods had been used after collection. One at a time, eighteen 50 ml falcon tubes were placed into the population cage for approximately one minute, so that flies could freely enter the tubes. Each tube contained around 20 to 40 flies of mixed sex. The eighteen falcon tubes were randomly allocated to one of the following six treatment groups, after which they were all frozen at -20°C:

1. Control – the tubes were immediately placed at -20°C.
2. Ethanol (99%) – 15 ml of ethanol was added to each tube.
3. Dry ice (2 h) – the tubes were immediately placed onto dry ice for 2 h.
4. Dry ice (24 h) – the tubes were immediately placed onto dry ice for 24 h.
5. Dry ice (2 h) + ethanol – the tubes were immediately placed into dry ice for 2 h then 15 ml of ethanol was added to each tube.
6. Dry ice (24 h) + ethanol – the tubes were immediately placed onto dry ice for 24 h then 15 ml of ethanol was added to each tube.

After one week we placed each fly on a microscope slide and placed a drop of Ringer's solution (182 mM KCl, 46 mM NaCl, 3 mM $CaCl_2.2H_2O$, 10 mM Tris-HCl) on top of it and examined it under a Leica M205C stereomicroscope for the presence/absence of bristles and antennae or damage to legs and wings.

In total 624 flies were examined for damage. Only two flies had missing leg parts and 39 had wing damage in this experiment, therefore the effects of treatment on these types of damage were not tested statistically. We therefore tested whether the storage method influenced the presence of head bristles, thorax bristles and antennae. Additionally, we tested the impact of sex on susceptibility to damage. Individual body parts were analysed separately with a generalized linear mixed model (glmer) with a

binomial distribution and falcon tube as a random factor. We included two-way interactions between treatment and sex to the model. A Type II ANOVA was applied to the results of the model to test the main effect of treatment and sex:
Supplementary Model 1: presence/absence of head bristles, thorax bristles or antennae ~ treatment × sex + (1 | falcon tube)

The storage methods had a significant effect on damage to various body parts: head bristles (LR Chisq = 54.82, df = 5, $p < 0.0001$), thorax bristles (LR Chisq = 55.18, df = 5, $p < 0.0001$), and antennae (LR Chisq = 17.3016, df = 3, $p < 0.0001$). After applying the correction for multiple testing, we found that treatments 4 (dry ice 24 h) and 6 (dry ice 24 h + ethanol) had a significant effect on the loss of bristles and antennae compared to the other treatments. Additionally, females exhibited more significant loss of thorax bristles compared to males (LR Chisq = 11.64, df = 1, $p < 0.0001$). As a result of this experiment, we chose to exclude the loss of bristles and antennae from the observations of wild collected flies, due to the possibility that these types of damage could be the result of the storage methods we had used. Due to the limited occurrence of leg and wing damage among the flies, we decided to conduct another experiment to reassess these specific types of damage.

***Experiment 2****: Testing the effect of storage methods on leg, wing and abdomen damage*
In a second experiment, we tested our methods again for missing leg parts and wing damage, as well as examining the flies for the presence/absence of melanized spots on the ventral abdomen. To determine if -20 °C by itself caused any damage, we included an additional control group that examined flies at room temperature immediately after being placed in the tubes. The experiment was done the same way as described for Experiment 1. Twenty-four falcon tubes were randomly allocated to one of the following the eight treatment groups and twenty females and twenty males were examined for each treatment:
1. Control 1 - Freezer – the tubes were immediately placed at -20 °C
2. Control 2 - Room temperature - the tubes were kept in the room temperature
3. Ethanol + freezer - 15 ml of ethanol was added to each tube and the tubes were immediately placed at -20 °C
4. Ethanol + room temperature - 15 ml of ethanol was added to each tube and the tubes were kept at the room temperature
5. Dry ice (24h) + freezer - the tubes were immediately placed into dry ice for 24h then placed at -20°C
6. Dry ice (24h) + ethanol + freezer - the tubes were immediately placed into dry ice for 24h then 15ml of ethanol was added to each tube and placed at -20°C
7. Dry ice (24h) + room temperature- the tubes were immediately placed into dry ice for 24h then were kept in the room temperature

8. Dry ice (24h) + ethanol + room temperature - the tubes were immediately placed into dry ice for 24h then 15ml of ethanol was added to each tube and were kept in the room temperature

Except for control 2 (room temperature), we examined flies under Ringer's solution, as described in Experiment 1. The flies in control 2 were still alive at the time of examination, therefore we examined them immediately under $CO_2$ and without Ringer's solution. The rest of the samples were examined one week after the treatment.

In total 960 flies were examined for damage. Only two flies had missing leg parts, therefore the effects of treatment on this type of damage were not tested statistically. To assess the effect of storage method on wing damage, a generalized linear model (glm) with a binomial distribution was used, with wing damage included as a binary response variable. We did not include the falcon tube into the model because the variance could not be calculated when we include this factor. A two-way interaction between treatment and sex was tested and a Type II ANOVA was applied to the results of the model to test the main effect of treatment and sex. The melanized spots on the body were tested with the same model as Supplementary Model 1 except sex was not included to the model since males did not exhibit this type of damage.

We observed a marginally significant effect of the storage method on wing damage (LR Chisq = 16.08, df = 7, $p = 0.024$, N = 960), but this effect did not remain significant after multiple testing. On the other hand, melanised spots were exclusively found in females, and the storage methods had a significant effect (LR Chisq = 16.08, df = 7, $p < 0.0001$, N = 306). After the multiple testing, the methods involving ethanol exhibited a significant effect on the presence of melanised spots. We interpreted this result as indicating that ethanol might enhance the visibility of melanised spots, leading to a higher number of females with these spots in this treatment. In light of these findings, we decided to include leg and wing damage in the main experiment, as we found no significant effect of our storage method on these types of damage. However, for the melanised spots on the ventral abdomen, we planned to conduct another experiment to specifically test the effect of ethanol on this type of damage.

***Experiment 3****: Testing the effect of ethanol on melanized spots on the female ventral abdomen*

In a third experiment, our aim was to further investigate whether ethanol had an enhancing effect on the visibility of melanised spots. We looked for melanized spots on the ventral abdomen and since only females have these spots, the experiment was carried out with only females the same way as described in Experiment 1. Twelve falcon tubes were randomly allocated to one of the following four treatment groups and were kept at room temperature. Ten to 20 females were examined for each treatment.

1. $CO_2$ - the flies were anesthetized with $CO_2$ and were examined without Ringer's solution

2. $CO_2$ + Ringer’s - the flies were anesthetized with $CO_2$ and were examined in Ringer’s solution
3. Ethanol - 15 ml of ethanol was added to each tube and then flies were examined without Ringer’s solution
4. Ethanol + Ringer’s - 15 ml of ethanol was added to each tube and then flies were examined in Ringer’s solution

In total 146 flies were examined in a random order immediately after being placed into a falcon tube. The effect of ethanol on melanized spots was tested using the same model as in Supplementary Model 1.

We observed a significant effect (LR Chisq = 11.59, df = 3, p = 0.0089, N = 146) of the storage method on the presence of melanised spots. However, after multiple testing, the ethanol + Ringer's treatment had marginally more melanized spots compared to the $CO_2$ treatment (z = 2.78, p = 0.028). Considering that the flies examined in the $CO_2$ treatment were not in Ringer's solution, we attributed this difference mainly to the difficulty in detecting these melanised spots when the flies are dry. Therefore, we retained the evaluation of these types of damage in the main experiment.

#### Supplementary Information 2

**Wounding in female *D. simulans***

*Statistical analyses*

Similar to *D. melanogaster*, for *D. simulans* females first we tested whether there is variation in the number of body parts that are damaged per fly and whether the body parts differed in their likelihood of being damaged. We performed Chi-square tests as described in 2.6.1. We tested whether combined damage HTLA was affected by season or site, using Model 1 (2.6.1). We then examined individual body parts separately, and tested whether season or site affected the number of flies with total abdomen damage and the location of abdomen damage (ventral or dorsal abdomen) using again Model 1. The head, legs, thorax, wings, and genital damage were not tested because of the low proportion of flies with damage to these body parts. We also tested whether the proportion of *D. simulans* females with encapsulated melanised parasitoid eggs, was affected by season or site.

*Results*

The flies differed significantly in the number of wounded body parts (Chi square = 41.28, df = 2, $p < 0.0001$, see Figure Supplementary Information 2). Flies most frequently showed wounds on one body part, with only a small percentage of the flies having two injuries, and no flies having more than two wounded body parts. Thirty six percent of the females showed at least one type of wound to the external cuticle (Figure Supplementary Information 2). The head, legs, thorax, and abdomen differed significantly in the frequency with which they were wounded (Chi square = 41.24, df = 3, $p < 0.0001$), with the abdomen most frequently showing damage.

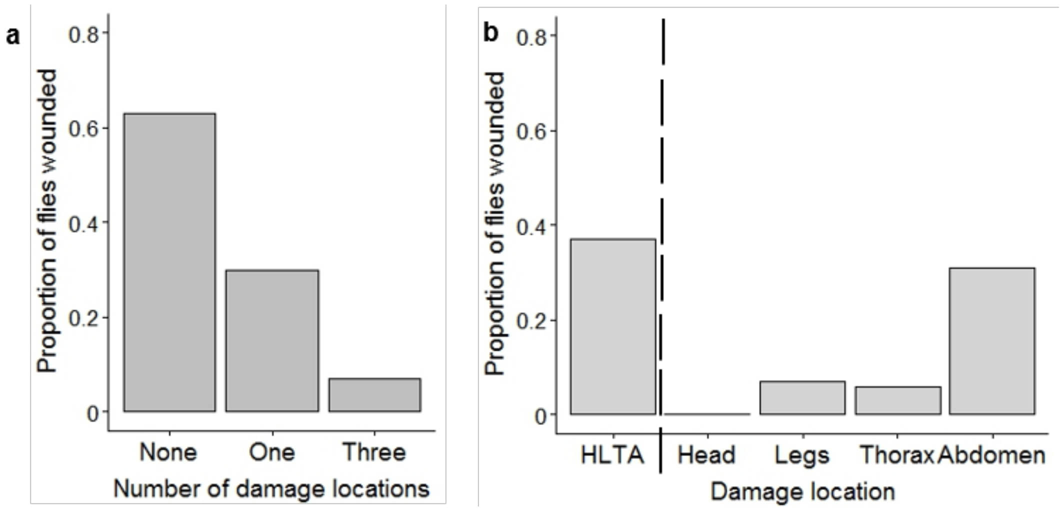


**Figure Supplementary Information 2. Frequency of wounding in *D. simulans*.** (a) The proportion of flies with none, one or more than one type of damage and (b) the proportion of wounding to the external cuticle (head, legs, thorax and abdomen).

When combining damage across the head, thorax, legs, and abdomen into one response variable, we observed a significant effect of site (LR Chisq = 11.07, df = 2, p = 0.0039). Post-hoc multiple comparisons revealed that *D. simulans* females from the D site had significantly more damage compared to the G (z = -3.116, p = 0.052) and L (z = -2.621, p = 0.0238) sites. Regarding total abdomen damage, we found significant effects of both season (LR Chisq = 9.02, df = 2, p = 0.0110) and site (LR Chisq = 12.91, df = 2, p = 0.0016). However, after post-hoc multiple comparisons, the effect of season was no longer significant. Still, the D site exhibited more abdominal damage than the L site (z = -2.96, p = 0.0088).

When considering abdomen damage based on whether it was to the ventral or dorsal abdomen, we found significant effects of both season (LR Chisq = 8.88, df = 2, p = 0.0118) and site (LR Chisq = 13.32, df = 2, p = 0.0013) on ventral abdomen damage. Post-hoc multiple comparisons showed that early summer flies had more damage compared to late summer flies (z = 2.46, p = 0.0375), and flies from the D site had more frequent damage to their ventral abdomen compared to the L site (z = -2.79, p = 0.0146). However, there was no significant effect of season and site on dorsal abdomen damage. Lastly, we did not find a significant effect of season or site on the presence of encapsulated parasitoid eggs.

#### Supplementary Tables

**Supplementary Table 1. The numbers of flies used to examine injuries per species, sex, season and collection site.** The species are: dmel = *D. melanogaster* and dsim = *D. simulans*; season: ES = Early summer, LS = Late summer and A = Autumn; and site: D = Domane Dahlem, G = Gartenbau and L = Lindicke. The same flies were used to check damage to the wing and the female genitalia, and these flies were a subset of the flies that were examined for combined damage to the head, thorax, legs, and abdomen (HTLA). N = sample size, NA = not applicable.

| *Species* | *Sex* | *Season* | *Site* | *N combined damage HTLA* | *N wing damage* | *N genital damage* |
|---|---|---|---|---|---|---|
| dmel | female | ES | D | 47 | 19 | 19 |
| dmel | female | ES | G | 32 | 20 | 20 |
| dmel | female | ES | L | 51 | 19 | 19 |
| dmel | female | LS | D | 85 | 19 | 19 |
| dmel | female | LS | G | 33 | 20 | 20 |
| dmel | female | LS | L | 93 | 20 | 20 |
| dmel | female | A | D | 35 | 17 | 17 |
| dmel | female | A | G | 87 | 18 | 16 |
| dmel | female | A | L | 74 | 13 | 13 |
| dmel | male | ES | D | 57 | 20 | NA |
| dmel | male | ES | G | 47 | 20 | NA |
| dmel | male | ES | L | 71 | 20 | NA |
| dmel | male | LS | D | 94 | 20 | NA |
| dmel | male | LS | G | 57 | 20 | NA |
| dmel | male | LS | L | 120 | 19 | NA |
| dmel | male | A | D | 35 | 20 | NA |
| dmel | male | A | G | 48 | 19 | NA |
| dmel | male | A | L | 95 | 15 | NA |
| dsim | female | ES | D | 2 | 1 | 1 |
| dsim | female | ES | G | 0 | 0 | 0 |
| dsim | female | ES | L | 1 | 1 | 1 |
| dsim | female | LS | D | 15 | 1 | 1 |
| dsim | female | LS | G | 0 | 0 | 0 |
| dsim | female | LS | L | 8 | 0 | 0 |
| dsim | female | A | D | 8 | 3 | 3 |
| dsim | female | A | G | 14 | 2 | 2 |
| dsim | female | A | L | 23 | 7 | 7 |

**Supplementary Table 3. The effects of sex, site, and season on combined damage HTLA (head, thorax, leg, and abdomen).** Statistically significant P-values are in bold font.

| *Tested effect* | *df* | *LR Chisq* | *P* |
|---|---|---|---|
| Sex | 1 | 22.07 | **< 0.0001** |
| Season | 2 | 6.47 | **0.0394** |
| Site | 2 | 1.57 | 0.46 |
| Sex × Season | 2 | 0.66 | 0.72 |
| Sex × Site | 2 | 2.18 | 0.34 |
| Season × Site | 4 | 21.82 | **< 0.0001** |

**Supplementary Table 4. The effects of sex, site, and season on thorax damage.** Statistically significant P-values are in bold font.

| *Tested effect* | *df* | *LR Chisq* | *P* |
|---|---|---|---|
| Sex | 1 | 2.97 | 0.08 |
| Season | 2 | 13.04 | **0.0015** |
| Site | 2 | 11.56 | **0.0031** |
| Sex × Season | 2 | 3.65 | 0.16 |
| Sex × Site | 2 | 0.30 | 0.86 |
| Season × Site | 4 | 5.75 | 0.22 |

**Supplementary Table 5. The effects of sex, site, and season on total, ventral and dorsal abdomen damage.** Statistically significant P-values are in bold font.

| *Response variable* | *Tested effect* | *df* | *LR Chisq* | *P* |
|---|---|---|---|---|
| Total abdomen damage | Sex | 1 | 32.60 | **< 0.0001** |
| | Season | 2 | 7.61 | **0.0223** |
| | Site | 2 | 4.44 | 0.11 |
| | Sex × Season | 2 | 0.31 | 0.86 |
| | Sex × Site | 2 | 1.20 | 0.55 |
| | Season × Site | 4 | 20.63 | **< 0.0001** |
| Ventral abdomen damage | Sex | 1 | 46.809 | **< 0.0001** |
| | Season | 2 | 11.616 | **0.0030** |
| | Site | 2 | 4.93 | 0.08 |
| | Sex × Season | 2 | 0.52 | 0.77 |
| | Sex × Site | 2 | 0.58 | 0.75 |
| | Season × Site | 4 | 21.26 | **< 0.0001** |
| Dorsal abdomen damage | Sex | 1 | 0.03 | 0.87 |
| | Season | 2 | 0.45 | 0.80 |
| | Site | 2 | 5.68 | 0.06 |
| | Sex × Season | 2 | 3.37 | 0.19 |
| | Sex × Site | 2 | 4.02 | 0.13 |
| | Season × Site | 4 | 3.27 | 0.51 |

**Supplementary Table 6. The effects of sex, site, and season on wing damage on veins and not on veins.** Statistically significant P-values are in bold font.

| *Response variable* | *Tested effect* | *df* | *LR Chisq* | *P* |
|---|---|---|---|---|
| Wing damage on veins | Sex | 1 | 0.45 | 0.50 |
| | Season | 2 | 10.42 | **0.0054** |
| | Site | 2 | 6.81 | **0.0332** |
| | Sex × Season | 2 | 1.96 | 0.37 |
| | Sex × Site | 2 | 4.88 | 0.09 |
| | Season × Site | 4 | 5.38 | 0.25 |
| Wing damage not on veins | Sex | 1 | 1.59 | 0.21 |
| | Season | 2 | 9.30 | **0.0096** |
| | Site | 2 | 4.42 | 0.11 |
| | Sex × Season | 2 | 0.77 | 0.68 |
| | Sex × Site | 2 | 1.44 | 0.49 |
| | Season × Site | 4 | 5.97 | 0.201 |

**Supplementary Table 7. The effects of sex, site, and season on genital damage.** Statistically significant P-values are in bold font.

| *Tested effect* | *df* | *LR Chisq* | *P* |
|---|---|---|---|
| Site | 1 | 16.19 | **< 0.0001** |
| Season | 2 | 1.29 | 0.52 |
| Site × Season | 2 | 8.28 | 0.082 |

**Supplementary Table 8. The effects of sex, site, and season on encapsulated parasitoid egg.** Statistically significant P-values are in bold font.

| *Tested effect* | *df* | *LR Chisq* | *P* |
|---|---|---|---|
| Sex | 1 | 3.35 | 0.07 |
| Season | 2 | 3.55 | 0.17 |
| Site | 2 | 6.57 | **0.0373** |
| Sex × Season | 2 | 0.88 | 0.64 |
| Sex × Site | 2 | 6.82 | **0.0330** |
| Season × Site | 4 | 17.34 | **0.0017** |

**Supplementary Table 9. The effects of sex, site, and season on probability of a mite infestation.** Statistically significant P-values are in bold font.

| *Tested effect* | *df* | *LR Chisq* | *P* |
|---|---|---|---|
| Sex | 1 | 1.74 | 0.19 |
| Season | 2 | 1.46 | 0.48 |
| Site | 2 | 4.83 | 0.09 |

#### Supplementary Figures

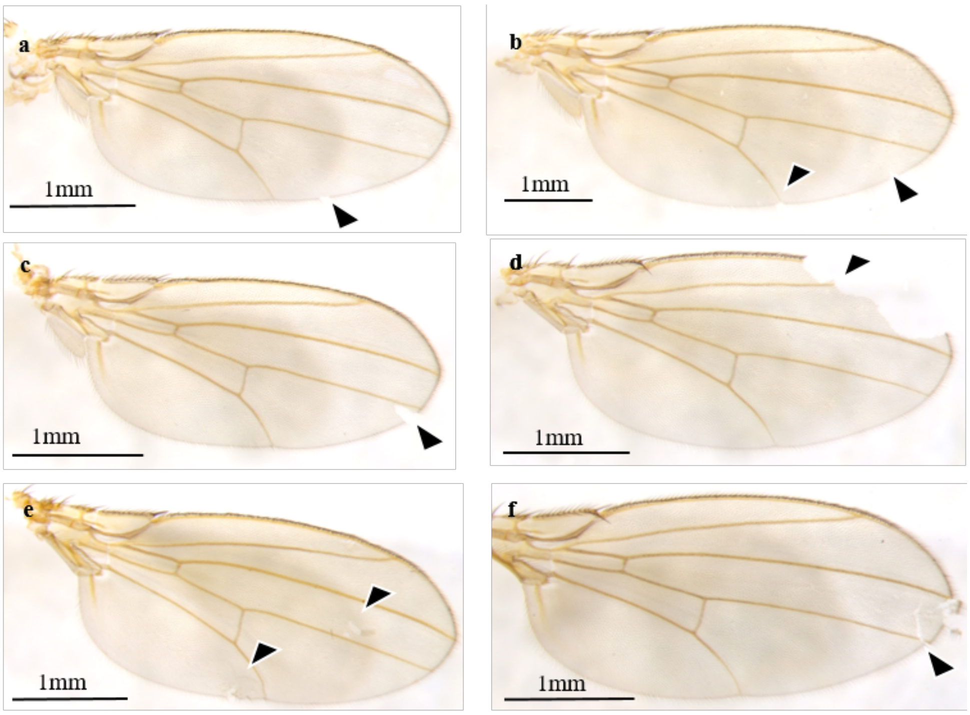


**Supplementary Figure 1. Examples of wing damage categories.** (a) notch not on vein, (b) notch on vein, (c) area not on vein, (d) area on vein, (e) tears not on vein and (f) tear on vein.

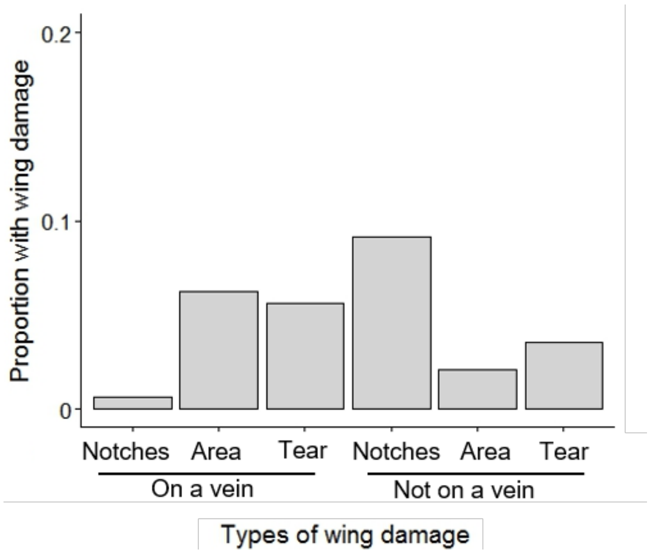


**Supplementary Figure 2. The prevalence of categories of wing damage in *D. melanogaster*.**
